# Generalization as diffusion: human function learning on graphs

**DOI:** 10.1101/538934

**Authors:** Charley M. Wu, Eric Schulz, Samuel J. Gershman

## Abstract

From social networks to public transportation, graph structures are a ubiquitous feature of life. How do humans learn functions on graphs, where relationships are defined by the connectivity structure? We adapt a Bayesian framework for function learning to graph structures, and propose that people perform generalization by assuming that the observed function values diffuse across the graph. We evaluate this model by asking participants to make predictions about passenger volume in a virtual subway network. The model captures both generalization and confidence judgments, and provides a quantitatively superior account relative to several heuristic models. Our work suggests that people exploit graph structure to make generalizations about functions in complex discrete spaces.

## Introduction

Most of function learning research has focused on how people learn a relationship between two continuous variables (Mcdaniel & Busemeyer, 2005; Lucas, Griffiths, Williams, & Kalish, 2015; DeLosh, Busemeyer, & McDaniel, 1997). How much hot sauce should I add to enhance my meal? How hard should I push a child on a swing? While function learning on continuous spaces is ubiquitous, many other relationships in the world are defined by functions on discrete spaces. For example, navigating a subway network and constructing a bookshelf both require representation of functions mapping discrete inputs (subway stops and configurations of components) to continuous outputs (passenger volume and probability of success). Likewise, language, commerce, and social networks are all defined partly by discrete relationships. How do people learn functions on discrete graph structures?

We propose that a diffusion kernel provides a suitable similarity metric based on the transition structure of a graph. When combined with the Gaussian Process (GP) regression framework, we arrive at a model of how humans learn functions and perform inference on graph structures. Using a virtual subway network prediction task, we pit this model against heuristic alternatives, which perform inference with lower computation demands, but are unable to capture human inference and confidence judgments. We also show that the diffusion kernel can be related to prominent models in continuous function learning and models of structure learning. This opens up a rich set of theoretical connections across theories of human learning and generalization.

### Computational Models of Function Learning

Based on a limited set of observations, how can you interpolate or extrapolate to predict unobserved data? This question has been the focus of human function learning research, which has traditionally studied predictions in continuous spaces (e.g., the relationship between two variables; Busemeyer, Byun, DeLosh, & McDaniel, 1997). Function learning research has revealed how inductive biases guide learning (Kwantes & Neal, 2006; Kalish, Griffiths, & Lewandowsky, 2007; Schulz, Tenenbaum, Duvenaud, Speekenbrink, & Gershman, 2017) and which types of functions are easier or harder to learn (Schulz, Tenenbaum, Reshef, Speekenbrink, & Gershman, 2015).

Several theories have been proposed to account for how humans learn functions. Earlier approaches used rule-based models that assumed a specific parametric family of functions (e.g., linear or exponential; Brehmer, 1974; Carroll, 1963; Koh & Meyer, 1991). However, the rigidity of rule-based learning struggled to account for order-of-difficulty effects in interpolation tasks (Mcdaniel & Busemeyer, 2005), and could not capture the biases displayed in extrapolation tasks (DeLosh et al., 1997).

An alternative approach relied on similarity-based learning, using connectionist networks to associate observed inputs and outputs (DeLosh et al., 1997; Kalish, Lewandowsky, & Kruschke, 2004; Mcdaniel & Busemeyer, 2005). The similarity-based approach is able to capture how people interpolate, but fails to account for some of the inductive biases displayed in extrapolation and in the partitioning of the input space. In some cases, hybrid architectures were developed to incorporate rule-based functions in a associative framework (e.g., Kalish et al., 2004; Mcdaniel & Busemeyer, 2005) in an attempt to gain the best of both worlds.

More recently, a theory of function learning based on GP regression was proposed to unite both accounts (Lucas et al., 2015), because of its inherent duality as both a rule-based and a similarity-based model. GP regression is a non-parametric method for performing Bayesian function learning (Schulz, Speekenbrink, & Krause, 2018), has successfully described human behavior across a range of traditional function learning paradigms (Lucas et al., 2015), and can account for compositional inductive biases (e.g., combining periodic and long range trends; Schulz et al., 2017).

While the majority of function learning research has studied continuous spaces, many real-world problems are discrete (Kemp & Tenenbaum, 2008). In a completely unstructured discrete space, the task of function learning is basically hopeless, because there is no basis for generalization across inputs. Fortunately, most real-world problems have structure (Tenenbaum, Kemp, Griffiths, & Goodman, 2011), which we can often represent as a connectivity graph that encodes how inputs (nodes) relate to each other (see Kemp & Tenenbaum, 2008, for a similar argument). By assuming that functions vary smoothly across the graph (a notion we formalize below), functions can be generalized to unobserved inputs. Although this idea has been studied extensively in machine learning, it has not yet fully permeated into studies of human function learning.

### Goals and Scope

We describe a model of learning graph-structured functions using a diffusion kernel. The diffusion kernel specifies the covariance between function values at different nodes of a graph based on its connectivity structure. When combined with the GP framework, it allows us to make Bayesian predictions about unobserved nodes. Even though previous work has investigated how people learn the relational structure of a graph (Kemp & Tenenbaum, 2008; Kemp, Tenenbaum, Griffiths, Yamada, & Ueda, 2006; Tomov, Yagati, Kumar, Yang, & Gershman, 2018), or infer properties of unobserved inputs (Kemp & Tenenbaum, 2009; Kemp, Shafto, & Tenenbaum, 2012), less is known about how people learn functions in discrete spaces with real-valued outputs.

We present an experiment where participants are shown a series of randomly generated subway maps and asked to predict the number of passengers at unobserved stations to test our model of function learning on graphs. In addition, we collected confidence judgments from participants. We compared the GP diffusion kernel model to heuristic models based on nearest-neighbor interpolation.

### Function Learning on Graphs

We can specify a graph 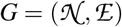 with nodes 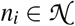 and edges 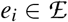 to represent a structured state space (Fig. 1a).

Nodes represent states and edges represent connections. For now, we assume that all edges are undirected (i.e., if *x → y* then *y → x*).

The diffusion kernel (Kondor & Lafferty, 2002) defines a similarity metric *k*(*s, s^!^*) between any two nodes on a graph based on the matrix exponentiation of the graph Laplacian:

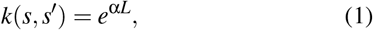

where *L* is the graph Laplacian:

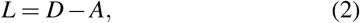

with the adjacency matrix *A* and the degree matrix *D*. Each element *a*_*i j*_ is 1 when nodes *i* and *j* are connected, and 0 otherwise, while the diagonals of *D* describe the number of connections of each node. The graph Laplacian can also describe graphs with weighted edges, where *D* becomes the weighted degree matrix and *A* becomes the weighted adjacency matrix.

Intuitively, the diffusion kernel assumes that function values diffuse along the edges similar to a heat diffusion process (i.e., the continuous limit of a random walk). The free parameter α governs the rate of diffusion, where α *→* 0 assumes complete independence between nodes, while α *→* ∞ assumes all nodes are perfectly correlated.

**Figure 1:**
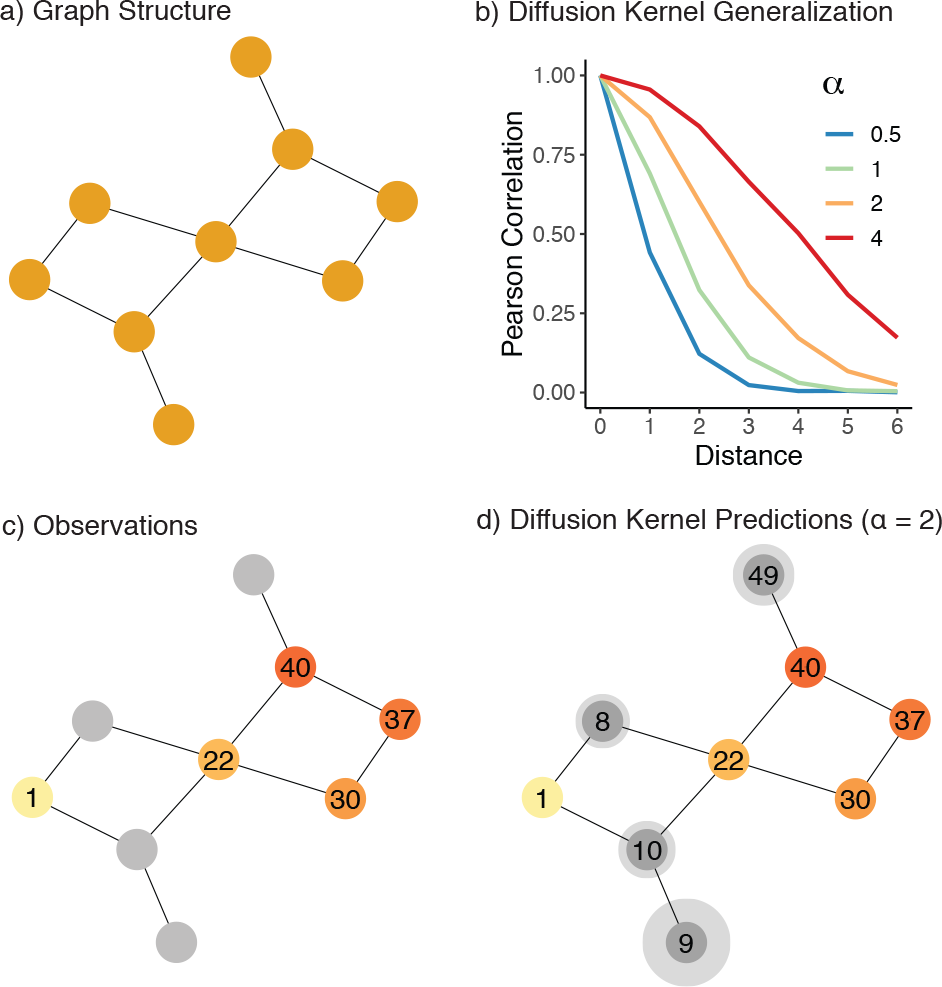
Graph-structured function learning. **a**) An example of a graph structure, where nodes represent states and edges indicate the transition structure. **b**) A diffusion kernel is a similarity metric between nodes on a graph, allowing us to generalize to unobserved nodes based on the assumption that the correlation between function values decays as an exponential function of the distance between two nodes. The diffusion parameter (α) governs the rate of decay. **c**) Given some observations on the graph (colored nodes), we can use the diffusion kernel combined with the Gaussian Process framework to make predictions (**d**) about expected rewards (numbers in grey nodes) and the underlying uncertainty (size of halo) for each unobserved node.

From the similarity metric defined by the diffusion kernel, we can use the GP regression framework (Rasmussen & Williams, 2006) to perform Bayesian inference over graph-structured functions. A GP defines a distribution over functions 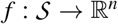 that map the input space 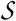 to real-valued scalar outputs:

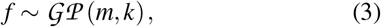

where *m*(*s*) is a mean function specifying the expected output of *s*, and *k*(*s, s^!^*) is the covariance function (kernel) that encodes prior assumptions about the smoothness of underlying function. Any finite set of function values drawn from a GP is multivariate Gaussian distributed.

We use the diffusion kernel (Eq. 1) to represent the co-variance *k*(*s, s^!^*) based on the connectivity structure of the graph, and follow the convention of setting the mean function to zero, such that the GP prior is fully defined by the kernel.

Given some observations 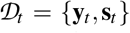 of observed out-puts **y**_*t*_ at states **s**_*t*_, we can compute the posterior distribution 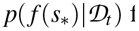 for any target state *s*_∗_. The posterior is a normal distribution with mean and variance defined as:

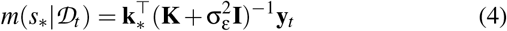

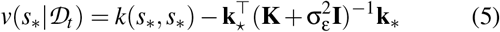

where **K** is the *t × t* covariance matrix evaluated at each pair of observed inputs, and **k**_∗_ = [*k*(*s*_1_, *s*_∗_), …, *k*(*s*_*t*_, *s*_∗_)] is the co-variance between each observed input and the target input *s*_∗_, and 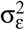 is the noise variance. Thus, for any node in the graph, we can make Bayesian predictions (Fig. 1e) about the expected function value 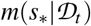 and also the level of uncertainty 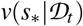.

The posterior mean function of a GP can be rewritten as:

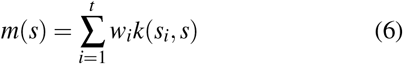

where each *s*_*i*_ is a previously observed state and the weights are collected in the vector 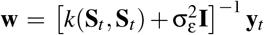. In-tuitively, this means that GP regression is equivalent to a linearly-weighted sum using basis functions *k*(*s*_*i*_, *s*) to project observed states onto a feature space (Schulz, Speekenbrink, & Krause, 2018). To generate new predictions for an unobserved state *s*, each output *y*_*t*_ is weighted by the similarity between observed states *s*_*t*_ and the target state *s*.

### Connections to Function Learning On Continuous Domains

The GP framework allows us to relate similarity-based function learning on graphs to theories of function learning in continuous domains. Consider the case of an infinitely fine lattice graph (i.e., a grid-like graph with equal connections for every node and with the number of nodes and connections approaching continuity). Following Kondor and Lafferty (2002) and using the diffusion kernel defined by Eq. 1, this limit can be expressed as

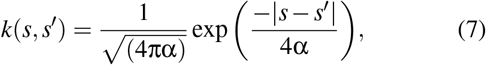

which is equivalent to a Radial Basis Function (RBF) kernel. Models similar to the RBF kernel are prominent in the literature on function learning in continuous domains (Busemeyer et al., 1997; Lucas et al., 2015). The RBF kernel has also been used to model how humans generalize about unobserved rewards in exploration tasks (Wu, Schulz, Speekenbrink, Nelson, & Meder, 2018). Thus, the RBF kernel can be understood as a special case of the diffusion kernel, when the underlying structure is symmetric and infinitely fine.

More broadly, both the RBF and diffusion kernel can be understood as instantiations of Shepard’s (1987) “universal law of generalization” in a function learning domain, by expressing generalization as an exponentially decaying function of the distance between two stimuli. Shepard famously proposed that the law of generalization should be the first law of psychology, while recent work has further entrenched it in fundamental properties of efficient coding (Sims, 2018) and measurement invariance (Frank, 2018).

### Heuristic Models

We compare the GP model to two heuristic strategies for function learning on graphs, which make predictions about the rewards of a target state *s*_∗_ based on a simple nearest neighbors averaging rule. The *k-Nearest Neighbors* (kNN) strategy averages the function values of the *k* closest states (including all states with same shortest path distance as the *k*-th closest), while the *d-Nearest Neighbors* (dNN) strategy averages the function values of all states within path distance *d*. Both kNN and dNN default to a prediction of 25 when the set of neighbors are empty (i.e., the median value in the experiment).

Both the dNN and kNN heuristics approximate the local structure of a correlated graph structure with the intuition that nearby states have similar function values. While they sometimes make the same predictions as the GP model and have lower computational demands, they fail to capture the connectivity structure of the graph and are unable to learn directional trends. Additionally, they only provide point-estimate predictions, and thus do not capture the underlying uncertainty of a prediction (which we use to model confidence judgments).

### Experiment: Subway Prediction Task

We used a “Subway Prediction Task” to study how people perform function learning in graph-structured state spaces. Participants were shown a series of graphs described as subway maps, where nodes corresponded to stations and edges indicated connections (Fig. 2). Participants were asked to predict the number of passengers (in a randomly selected train car) at a target station, based on observations from other stations.

### Methods and procedure

We recruited 100 participants (*M*_*age*_ = 32.7; *SD* = 8.4; 28 female) on Amazon MTurk to perform 30 rounds of a graph prediction task. On each graph, numerical information was provided about the number of passengers at 3, 5, or 7 other stations (along with a color aid), from which participants were asked to predict the number of passengers at a target station and provide a confidence judgment (Likert scale from 1-11). The subway passenger cover story was used to provide intuitions about graph correlated functions. Additionally, participants observed 10 fully revealed graphs to familiarize themselves with the task and completed a comprehension check before starting the task. Participants were paid a base fee of $2.00 USD for participation with an additional performance contingent bonus of up to $3.00 USD. The bonus payment was based on the mean absolute judgment error weighted by confidence judgments: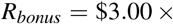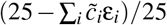 where 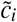 is the normalized confidence judgment 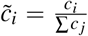 and ε_*i*_ is the absolute error for judgment *i*. On average, participants completed the task in 8.09 minutes (*SD* = 3.7) and earned $3.87 USD (*SD* = $0.33).

**Figure 2:**
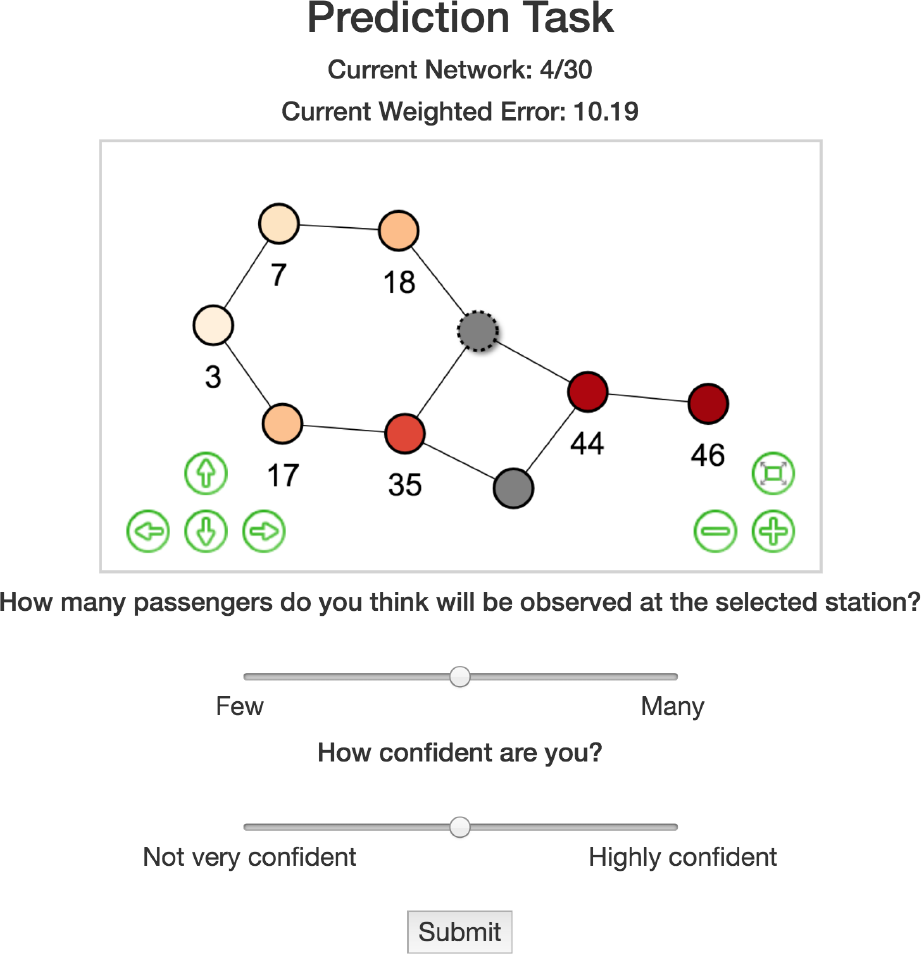
Screenshot from the Subway Prediction Experiment. Observed nodes (3, 5, or 7 randomly sampled nodes depending on the information condition) are shown with a numerical value and a corresponding color aid (darker indicates larger values). The target node is indicated by the dashed line, and dynamically changes color and displays a numerical value when participants move the top slider. Confidence judgments were used to compute a weighted error (i.e., more confident answers having a larger contribution), which was used to determine the performance contingent bonus.

In each of the 30 rounds, a different graph was sampled without replacement. We used three different information conditions (observations ∈ [3, 5, 7]; each used in 10 rounds in randomly shuffled order) as a within-subject manipulation determining the number of randomly sampled nodes with revealed information. In each round, participants were asked to predict the value of a target node, which was randomly sampled from the remaining unobserved nodes.

All participants observed the same set of 40 graphs that were sampled without replacement for the 10 fully revealed examples in the familiarization phase and for the 30 graphs in the prediction task. We generated the set of 40 graphs by iteratively building 3 *×* 3 lattice graphs (also known as mesh or grid graphs), and then randomly pruning 2 out of the 12 edges. In order to generate the functions (i.e., number of passengers), we fit a diffusion kernel to the graph and then sampled a single function from a GP prior, where the diffusion parameter was set to α = 2.

## Results

Figure 3 shows the behavioral and model-based results of the experiment. We applied linear mixed-effects regression to estimate the effect of the number of observed nodes on participant prediction errors, with participants as a random effect. Participants made systematically lower error predictions as the number of observable nodes increased (β = −.11, *t*(99) = −6.28, *p* < .001, *BF* > 100^1^; Fig. 3a). Repeating the same analysis but using participant confidence judgments as the dependent variable, we found that confidence increased with the number of observable nodes (β = .16, *t*(99) = 11.3, *p* < .001, *BF* > 100; Fig. 3b). Finally, participants were also able to calibrate confidence judgments to the accuracy of their predictions, with higher confidence predictions having consistently lower error (β = −.19, *t*(99) = −9.0, *p* < .001, *BF* > 100; Fig. 3c). There were no substantial effects of learning over rounds (β = .01, *t*(99) = 0.47, *p* = .642, *BF* = 0.2), suggesting the familiarization phase and cover story were sufficient for providing intuitions about graph correlated structures.

### Model comparison

We compare the predictive performance of the GP with the dNN and kNN heuristic models. Using participant-wise leave-one-out cross-validation, we estimate model parameters for all but one judgment, and then make out-of-sample predictions for the left-out judgment. We repeat this procedure for all trials and compare predictive performance using Root Mean Squared Error (RMSE) over all left-out trials.

Figure 3d shows that the GP made better predictions than both the dNN (*t*(99) = −4.06, *p* < .001, *d* = 0.41, *BF* > 100) and kNN models (*t*(99) = −7.19, *p* < .001, *d* = 0.72, *BF* > 100). Overall, 58 out of 100 participants were best predicted by the GP, 31 by the dNN, and 11 by the kNN. Figure 3e shows individual parameter estimates of each model. The estimated diffusion parameter α was not substantially different from the ground truth of α = 2 (*t*(99) = −0.66, *p* = .51, *d* = 0.07, *BF* = 0.14), although the distribution appeared to be bimodal, with participants often underestimating or over-estimating the correlational structure. Estimates for *d* and *k* were highly clustered around the lower limit of 1, suggesting that averaging over larger portions of the graph were not consistent with participant predictions.

Finally, an advantage of the GP is that it produces Bayesian uncertainty estimates for each prediction. While the dNN and kNN models make no predictions about confidence, the GP uncertainty estimates correspond to participant confidence judgments (β = −.10, *t*(99) = −3.39, *p* < .001, *BF* > 100; linear mixed-effects model with participant as a random effect).

**Figure 3:**
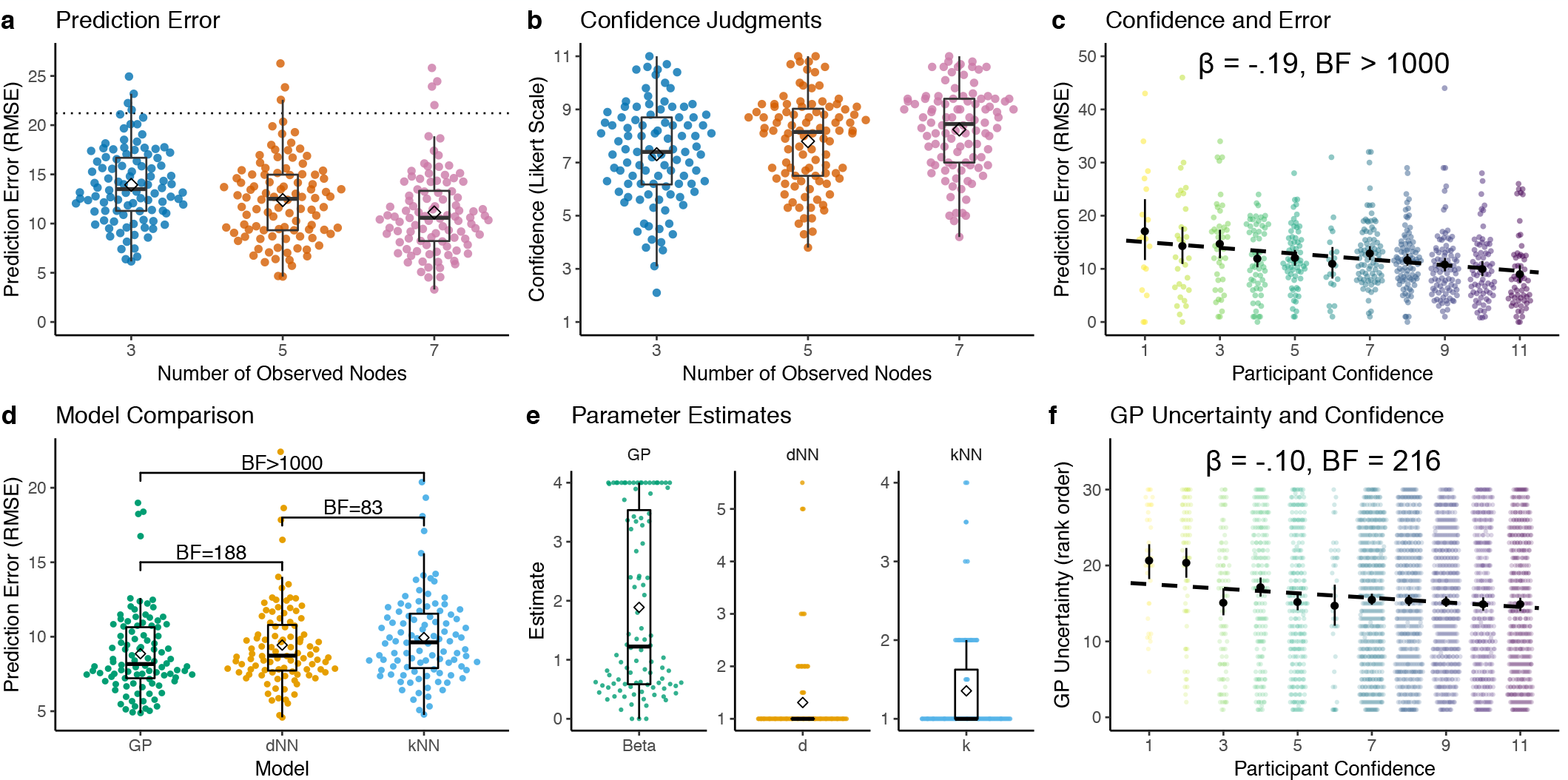
Results. **a-b**) Participant judgment errors and confidence estimates. Each dot is a single participant (averaged over each number of observed nodes), with Tukey boxplots and diamonds indicating group means. The dotted line in **a**) is a random baseline. **c**) Judgment error and confidence. Each colored dot is a participant (averaged over each confidence level), dashed line is a linear regression, with black dots and error bars indicating group means and 95% CI. We report the mixed-effects regression coefficient and Bayes Factor above. **d**) Cross-validated model comparison between the Gaussian Process with diffusion kernel (GP), d-nearest neighbors (dNN), and k-nearest neighbors (kNN). Each point is a single participant with a Tukey boxplot overlaid and diamonds indicating group means. Comparisons are for a Bayesian one-sample *t*-test, where the null hypothesis posits no difference between models and assumes a Cauchy prior with the scale set to 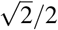. **e**) Parameter estimates, where each dot is the mean cross-validated estimate for each participant, with Tukey boxplots and diamonds indicating group means. **f**) GP uncertainty estimates (rank ordered within participant) and participant confidence judgments (Likert scale). Dotted line is a linear regression, with black dots and error bars indicating mean and 95% CI. We report the mixed-effects regression coefficient and Bayes factor (see text for details).

## Discussion

How do people learn about functions on structured discrete spaces like graphs? We show how a GP with a diffusion kernel can be used as a model of function learning that produces Bayesian predictions about unobserved nodes. Our model integrates existing theories of human function learning in continuous spaces, where the RBF kernel (commonly used in continuous domains) can be seen as a special limiting case of the diffusion kernel. Using a virtual subway task, we show that the GP was able to capture how people make judgments about unobserved nodes and is also able to generate uncertainty estimates that correspond to participant confidence ratings.

### Related work

Previous work has also investigated how people perform inference over graphs (Kemp & Tenenbaum, 2009, 2008; Shafto, Kemp, Baraff, Coley, & Tenenbaum, 2005; Tomov et al., 2018). Whereas these studies were geared towards probing how people inferred underlying structure (Kemp & Tenenbaum, 2008) and how (implicit or explicit) representations of structure influenced causal property judgments (Kemp & Tenenbaum, 2009; Shafto et al., 2005), the goal of our Subway Prediction Task was to study how people perform functional inference given explicit knowledge of a relational structure. Thus, our study can be seen as a real-valued extension of the experiments presented in Kemp and Tenenbaum (2009) and Shafto et al. (2005), where we explicitly present the underlying structure and modeled both participants predictions and their confidence judgments simultaneously.

Our approach also has formal similarities to Kemp and Tenenbaum (2008, 2009), who used a kernel defined as 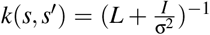 to generate feature vectors over structured representations, in order to approximate a prior over properties distributed across the graph. This kernel is a reformulation of the regularized Laplacian kernel^2^ (Zhu, Lafferty, & Ghahramani, 2003), which belongs to the same broad framework of regularization operators (Smola & Kondor, 2003) as the diffusion kernel (Eq. 1), with both providing similar inductive biases of smoothness over the graph struc ture.

### Future Work and Limitations

Currently, we have only studied how people learn functions on spatial representations of graph structures, where all nodes and edges are visible simultaneously. However, people can perform inferences over discrete structures that are more conceptual such as social hierarchies (Lau, Pouncy, Gershman, & Cikara, 2018) or causal connections (Rothe, Deverett, Mayrhofer, & Kemp, 2018). Given that the GP framework can be used to compare how people learn functions over different (i.e., spatial and conceptual) domains (Wu, Schulz, Garvert, Meder, & Schuck, 2018), comparing functional inference over conceptual and spatial graphs seems like promising extension for future studies.

Additionally, one could also assess the suitability of the diffusion kernel as a model for more complex problems, such as multi-armed bandit tasks with structured rewards (e.g., Schulz, Franklin, & Gershman, 2018) and in planning problems, where exploration plays a fundamental role. One advantage of the GP diffusion kernel model is that it makes prediction with estimates of the underlying uncertainty. Whereas many models of generalization only make point-estimates about the value of a state, the GP framework offers opportunities for using uncertainty-guided exploration strategies (e.g., Auer, 2002).

One limitation of the diffusion kernel is that it assumes *a priori* knowledge of the graph structure. While this may be a reasonable assumption in problems such as navigating a subway network where one can simply look at a map, this is not always the case. In contrast, the SR can learn the graph structure through experience (using prediction-error updating). Thus, the connection between the SR and the diffusion kernel presents a promising avenue for incorporating a plausible process model of structure learning.

## Conclusion

We show that GP regression, together with a diffusion kernel, captures how participants learn functions and make confidence ratings on graph structures in a virtual subway prediction task. Our model opens up a rich set of theoretical connections to theories of function learning on continuous domains and models of structure learning and property induction.

## Acknowledgements

CMW is supported by the International Max Planck Research School on Adapting Behavior in a Fundamentally Uncertain World; ES is supported by the Harvard Data Science Initiative

## Footnotes

1 β is the standardized effect size ∈ [−1, 1] and we approximate the Bayes Factor using bridge sampling (Gronau, Singmann, & Wagenmakers, 2017) to compare our model to an alternative intercept only null model, where both models were hierarchical regressions but only the alternative model contained the variable of interest as a regressor.

2 *k*(*s, s^!^*) = (*I* + σ^2^*L*)^−1^

## References

Auer, P. (2002). Using confidence bounds for exploitation-exploration trade-offs. Journal of Machine Learning Research, 3, 397–422.

Brehmer, B. (1974). Hypotheses about relations between scaled variables in the learning of probabilistic inference tasks. Organizational Behavior and Human Performance, 11(1), 1–27.

Busemeyer, J. R., Byun, E., DeLosh, E. L., & McDaniel, M. A. (1997). Learning functional relations based on experience with input-output pairs by humans and artificial neural networks. In K. Lamberts & D. Shanks (Eds.), Concepts and Categories (p. 405–437). Cambridge: MIT Press.

Carroll, J. D. (1963). Functional learning: The learning of continuous functional mappings relating stimulus and response continua. ETS Research Bulletin Series, 1963(2), i–144.

DeLosh, E. L., Busemeyer, J. R., & McDaniel, M. A. (1997). Extrapolation: The sine qua non for abstraction in function learning. Journal of Experimental Psychology: Learning, Memory, and Cognition, 23(4), 968.

Frank, S. A. (2018). Measurement invariance explains the universal law of generalization for psychological perception. Proceedings of the National Academy of Sciences, 115(39), 9803–9806. doi: 10.1073/pnas.1809787115

Gronau, Q. F., Singmann, H., & Wagenmakers, E.-J. (2017). Bridge-sampling: An R package for estimating normalizing constants. arXiv preprint arXiv:1710.08162.

Kalish, M. L., Griffiths, T. L., & Lewandowsky, S. (2007). Iterated learning: Intergenerational knowledge transmission reveals inductive biases. Psychonomic Bulletin & Review, 14(2), 288–294.

Kalish, M. L., Lewandowsky, S., & Kruschke, J. K. (2004). Population of linear experts: knowledge partitioning and function learning. Psychological Review, 111(4), 1072.

Kemp, C., Shafto, P., & Tenenbaum, J. B. (2012). An integrated account of generalization across objects and features. Cognitive Psychology, 64(1-2), 35–73.

Kemp, C., & Tenenbaum, J. B. (2008). The discovery of structural form. Proceedings of the National Academy of Sciences, 105(31), 10687–10692.

Kemp, C., & Tenenbaum, J. B. (2009). Structured statistical models of inductive reasoning. Psychological Review, 116(1), 20–58.

Kemp, C., Tenenbaum, J. B., Griffiths, T. L., Yamada, T., & Ueda, N. (2006). Learning systems of concepts with an infinite relational model. In AAAI (Vol. 3, p. 5).

Koh, K., & Meyer, D. E. (1991). Function learning: Induction of continuous stimulus-response relations. Journal of Experimental Psychology: Learning, Memory, and Cognition, 17(5), 811–836.

Kondor, R. I., & Lafferty, J. (2002). Diffusion kernels on graphs and other discrete structures. In Proceedings of the 19th International Conference on Machine Learning (Vol. 2002, pp. 315–322).

Kwantes, P. J., & Neal, A. (2006). Why people underestimate y when extrapolating in linear functions. Journal of Experimental Psychology: Learning, Memory, and Cognition, 32(5), 1019–1030.

Lau, T., Pouncy, H. T., Gershman, S. J., & Cikara, M. (2018). Discovering social groups via latent structure learning. Journal of Experimental Psychology: General, 147(12), 1881–1891.

Lucas, C. G., Griffiths, T. L., Williams, J. J., & Kalish, M. L. (2015). A rational model of function learning. Psychonomic Bulletin & Review, 22, 1193–1215.

Mcdaniel, M. A., & Busemeyer, J. R. (2005). The conceptual basis of function learning and extrapolation: Comparison of rule-based and associative-based models. Psychonomic Bulletin & Review, 12(1), 24–42.

Rasmussen, C., & Williams, C. (2006). Gaussian Processes for Machine Learning. MIT Press.

Rothe, A., Deverett, B., Mayrhofer, R., & Kemp, C. (2018). Successful structure learning from observational data. Cognition, 179, 266–297.

Schulz, E., Franklin, N. T., & Gershman, S. J. (2018). Finding structure in multi-armed bandits. bioRxiv, 432534.

Schulz, E., Speekenbrink, M., & Krause, A. (2018). A tutorial on gaussian process regression: Modelling, exploring, and exploiting functions. Journal of Mathematical Psychology, 85, 1–16.

Schulz, E., Tenenbaum, J. B., Duvenaud, D., Speekenbrink, M., & Gershman, S. J. (2017). Compositional inductive biases in function learning. Cognitive Psychology, 99, 44–79.

Schulz, E., Tenenbaum, J. B., Reshef, D. N., Speekenbrink, M., & Gershman, S. (2015). Assessing the perceived predictability of functions. In Proceedings of the 37th Annual Meeting of the Cognitive Science Society (p. 2116–2121). Cognitive Science Society.

Shafto, P., Kemp, C., Baraff, E., Coley, J., & Tenenbaum, J. (2005). Context-sensitive induction. In Proceedings of the 27th Annua Conference of the Cognitive Science Society (pp. 2003–2008). Austin, TX: Cognitive Science Society.

Shepard, R. N. (1987). Toward a universal law of generalization for psychological science. Science, 237(4820), 1317–1323.

Sims, C. R. (2018). Efficient coding explains the universal law of generalization in human perception. Science, 360(6389), 652–656.

Smola, A. J., & Kondor, R. (2003). Kernels and regularization on graphs. In Learning theory and kernel machines (pp. 144–158). Springer.

Tenenbaum, J. B., Kemp, C., Griffiths, T. L., & Goodman, N. D. (2011). How to grow a mind: Statistics, structure, and abstraction. Science, 331(6022), 1279–1285.

Tomov, M., Yagati, S., Kumar, A., Yang, W., & Gershman, S. (2018). Discovery of hierarchical representations for efficient planning. bioRxiv, 499418.

Wu, C. M., Schulz, E., Garvert, M. M., Meder, B., & Schuck, N. W. (2018). Connecting conceptual and spatial search via a model of generalization. In T. T. Rogers, M. Rau, X. Zhu, & C. W. Kalish (Eds.), Proceedings of the 40th annual conference of the cognitive science society (pp. 1183–1188). Austin, TX: Cognitive Science Society.

Wu, C. M., Schulz, E., Speekenbrink, M., Nelson, J. D., & Meder, B. (2018). Generalization guides human exploration in vast decision spaces. Nature Human Behaviour, 2, 915–924. doi: 10.1038/s41562-018-0467-4

Zhu, X., Lafferty, J. D., & Ghahramani, Z. (2003). *Semi-supervised learning: From gaussian fields to gaussian processes* (Tech. Rep. No. CMU-CS-03-175). Carnegie Mellon University.

